# A multi-domain connector links the outer membrane and cell wall in deep-branching bacteria

**DOI:** 10.1101/2022.05.18.492506

**Authors:** Andriko von Kügelgen, Sofie van Dorst, Vikram Alva, Tanmay A. M. Bharat

## Abstract

*Deinococcus radiodurans* is a deep-branching extremophilic bacterium that is remarkably tolerant to numerous environmental stresses, including large doses of ultraviolet radiation and extreme temperatures. It can even survive in outer space for several years. This endurance of *D. radiodurans* has been partly ascribed to its atypical cell envelope comprising an inner membrane, a large periplasmic space with a thick peptidoglycan (PG) layer, and an outer membrane (OM) covered by a surface layer (S-layer). Despite intense research, molecular principles governing envelope organization and OM stabilization are unclear in *D. radiodurans* and related bacteria. Here, we report an electron cryomicroscopy (cryo-EM) structure of the abundant *D. radiodurans* OM protein SlpA, showing how its C-terminal segment forms homotrimers of 30-stranded β-barrels in the OM, whereas its N-terminal segment forms long, homotrimeric coiled coils linking the OM to the PG layer via S-layer homology (SLH) domains. Using the power of structure prediction and sequence-based bioinformatics, we further show that SlpA-like proteins are widespread in deep-branching Gram-negative bacteria, plausibly constituting an ancestral superfamily of OM-PG connectors, important for organizing the cell envelopes of many bacteria. Finally, combining our atomic structures with tomography of cell envelopes, we report a model for the cell surface of *D. radiodurans*, with implications on understanding the cell surface organization and hyperstability of *D. radiodurans* and related bacteria. Furthermore, the widespread occurrence of SlpA-like OM-PG connectors in deep-branching bacteria will help in understanding the evolutionary transition between Gram-negative and Gram-positive bacteria.

## Main text

*Deinococcus radiodurans* is an evolutionarily deep-branching bacterium with several distinctive characteristics (1). It is remarkably tolerant to large doses of ionizing and ultraviolet radiation, extreme temperatures, osmotic pressure, oxidative stress, and desiccation, primarily owing to its extensive DNA repair system (2), complex cell envelope (3), and antioxidation systems, such as the one involving the carotenoid deinoxanthin (4, 5). In fact, *D. radiodurans* can even survive for several years in outer space (6). Due to its ability to survive under extreme environmental conditions and its deep position in the bacterial tree of life, *D. radiodurans* has been of tremendous interest for several synthetic biology and evolutionary studies (2).

The cell envelope of *D. radiodurans* is atypical. While it stains Gram-positive, its architecture resembles that of Gram-negative bacteria, containing an inner membrane (IM) covered by a PG layer in a large periplasmic space (7–9) and an OM. However, unlike typical Gram-negative bacterial OMs, this OM lacks lipopolysaccharide and phospholipids, and instead has a lipid composition similar to the IM (10). Additionally, the *D. radiodurans* OM is covered by a regularly-spaced, hexagonal S-layer (11, 12). Previous studies have suggested that the S-layer is made of a protein called <u>H</u>exagonally <u>P</u>acked Interlayer (HPI) (3, 8, 11, 13–17), while newer studies have suggested that a hetero-complex with gating properties, termed the S-layer deinoxanthin-binding complex (SDBC), forms a large part of the *D. radiodurans* cell envelope including the S-layer (18, 19). A previously identified abundant protein called SlpA (UniProtKB Q9RRB6) is suggested to be the main component of this complex. Recently, an 11 Å resolution structure of this complex was reported using electron cryomicroscopy (cryo-EM), showing how it exhibits a triangular base that is partly embedded in the OM and a stalk departing orthogonally from the base, presumably away from the membrane (18).

In addition to the biophysical observations introduced above, from an evolutionary perspective, an ortholog of *D. radiodurans* SlpA (UniProtKB Q5SH37) has also been charaterised from the closely related thermophilic model bacterium *Thermus thermophilus* (20, 21). In both these organisms, deletion or truncation of *slpA* leads to remarkable disruption of the cell envelope (22, 23), underpinning its importance in cell surface organization. At the sequence level, SlpA contains a signal peptide, an SLH domain, a long, predicted α-helical region, and a C-terminal β-strand rich domain, which is thought to fold into an OM β-barrel (18, 19) (Figure 1A). Due to the presence of the N-terminal SLH domain, which commonly attaches S-layer proteins (SLPs) (21, 24–27) of Gram-positive bacteria to PG-linked pyruvylated secondary cell wall polymers (SCWPs), it has been suggested that SlpA constitutes the S-layer. Conversely, in *T. thermophilus*, SlpA has been shown to interact with PG through its SLH domain, suggesting a role for it as a periplasmic spacer (28). The role of SlpA in organising the cell envelope of *D. radiodurans* and related deep-branching bacteria such as *T. thermophilus* is, therefore, still enigmatic.

**Figure 1.**
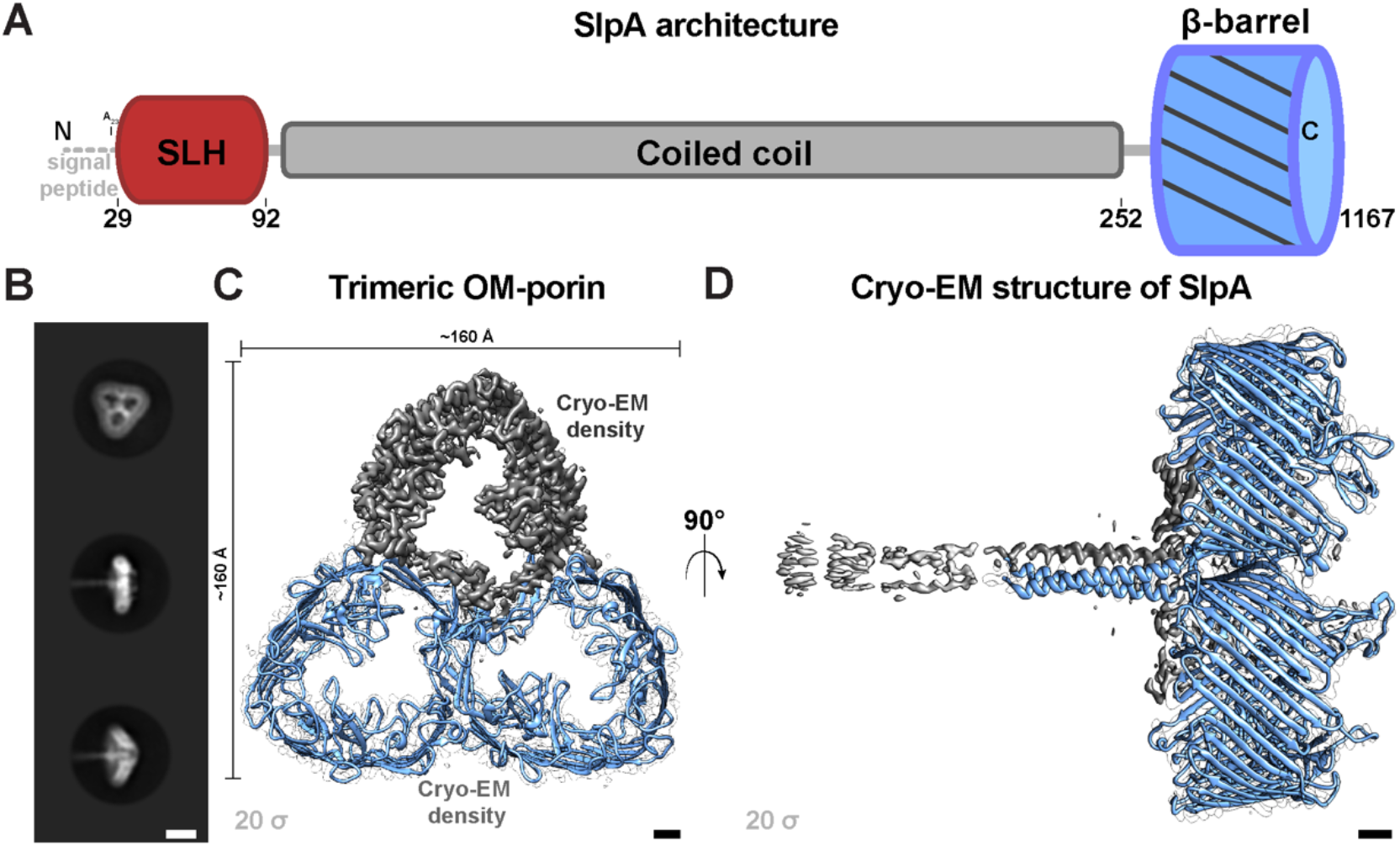
Cryo-EM reconstruction of *D. radiodurans* SlpA at 3.3 Å resolution. (A) The SlpA protein contains a tripartite structure including an N-terminal SLH domain which is connected to a C-terminal β-barrel by a long coiled-coil segment. (B) Two-dimensional class averages of the trimeric SlpA specimen used for cryo-EM structure determination. Characteristic top and side views are shown. (C) Density map of the SlpA trimer (contour level on the lower left of panel) shown from the top. Two subunits are shown as blue ribbons inside white envelope outlines and one as grey density (model hidden). (D) An orthogonal view of panel C), with the SlpA trimer shown from side. The extended coiled coil degrades in resolution towards the N-terminus (see also **Figure S1**), presumably due to flexibility of the long stalk. Scale bars: B) 100 Å; C-D) 10 Å

In this study, we report the electron cryomicroscopy (cryo-EM) structure of the SlpA protein complex from *D. radiodurans.* Our structure shows that SlpA exhibits a tripartite organization, with its C-terminal part forming a homotrimeric 30-stranded OM β-barrel, its central part forming a trimeric coiled coil that can traverse the large periplasmic space, and the extreme N-terminal part forming an SLH domain trimer that can interact with the PG layer. Our structure-and sequence-based bioinformatic analyses further show the presence of similar proteins in several phyla of deep-branching Gram-negative bacteria. Finally, combining our atomic structures and bioinformatic results with tomography of cell envelopes, we report a model for the cell envelope of *D. radiodurans,* showing how this Gram-negative (diderm) bacterial SlpA protein shares several characteristics commonly found in Gram-positive (monoderm) SLPs, with connotations on prokaryotic evolution.

## Results

### Overall structure of the *D. radiodurans* SlpA complex

To understand the molecular details of SlpA, we utilized previously described techniques (18, 19) to purify SlpA from *D. radiodurans* using detergent solubilization (Figure S1, Methods). Cryo-EM images of the purified specimen showed single-particles on the grid (Figure S1), which appeared to be made up of trimeric densities (Figure 1B), as reported previously (18). We performed single particle analysis on this cryo-EM data to solve a global 3.3 Å resolution structure of SlpA (Figures 1C-D, S1 and Table S1). The structure showed that SlpA forms a homotrimer of 30-antiparallel-stranded β-barrels (30 β-strands per SlpA monomer). The SlpA β-barrel is the first structurally characterized 30-stranded barrel and one of the largest single-chain β-barrels observed (29, 30). Since the SlpA complex was stabilized in detergent, and because the SlpA protein sequence possesses a β-signal motif, which is important for efficient targeting of OM β-barrel proteins (OMBBs) to the β-barrel assembly machine (BAM) complex (Figure S2) (31), it is highly likely that the β-barrel is present in the OM of *D. radiodurans* (Figure S2), in line with previous results on *slpA* deletion mutants in *D. radiodurans* (22).

Bioinformatic analyses revealed that homologs of SlpA are widespread in the Deinococcus-Thermus phylum, with some organisms, such as *Deinococcus wulumuqiensis* and *T. thermophilus,* even possessing two copies of SlpA (Table S2). The OMBB domain represents the most divergent part of SlpA proteins and contains either 28 or 30 β-strands depending on the species (Figure S3 and Table S2). For example, while the SlpA OMBB of *D. radiodurans, D. wulumuqiensis, Oceanithermus desulfurans, and Marinithermus hydrothermalis* possesses 30 strands, the SlpA OMBB of *T. thermophilus, Meiothermus ruber, and Deinococcus ficus* possesses 28-strands (Table S2). The C-terminal OMBB of the *D. radiodurans* SlpA is preceded by a long, homotrimeric coiled-coil segment, which, in our cryo-EM map, is well-resolved from residue 215 onwards. Together, there are extensive protein:protein homotrimeric interfaces both in the β-barrel and in the coiled-coil segment that appear to stabilize the trimeric SlpA complex (Figure 1C-D).

### SlpA is arranged as a blocked β-barrel with a coiled-coil stalk connecting the OM to PG

When compared to its homologs, the OMBB of *D. radiodurans* SlpA (residues 254-1167) contains several insertions that block the pore (Figures 2A and S3). Residues 272-377 form the most extensive, ordered insertion that lines the cavity of the pore. This insertion appears to be stabilized by a canonical bacterial SLP metal-ion binding site (32) coordinated by residues D274, D276, and D310 (Figure S4). Likewise, large insertions blocking the pore, with putative metal-ion binding sites, are found between residues 429-471 and 686-753 (Figures 2B and S4). Sequence analysis reveals that these insertions are only present in closely related Deinococcales (e.g., *D. wulumuqiensis*) and are less extensive or absent in SlpA proteins of most other Deinococcales and Thermales (Figure S3), suggesting that SlpA of *D. radiodurans* may not fulfil a role as a pore, but rather functions as an abundant membrane scaffold organizing the cell envelope. Also, since many bacteria in the Deinococcus-Thermus phylum do not possess an S-layer built of the HPI protein, the extensive insertions in *D. radiodurans* may also be involved in anchoring the HPI-based S-layer, as previously suggested (18).

**Figure 2.**
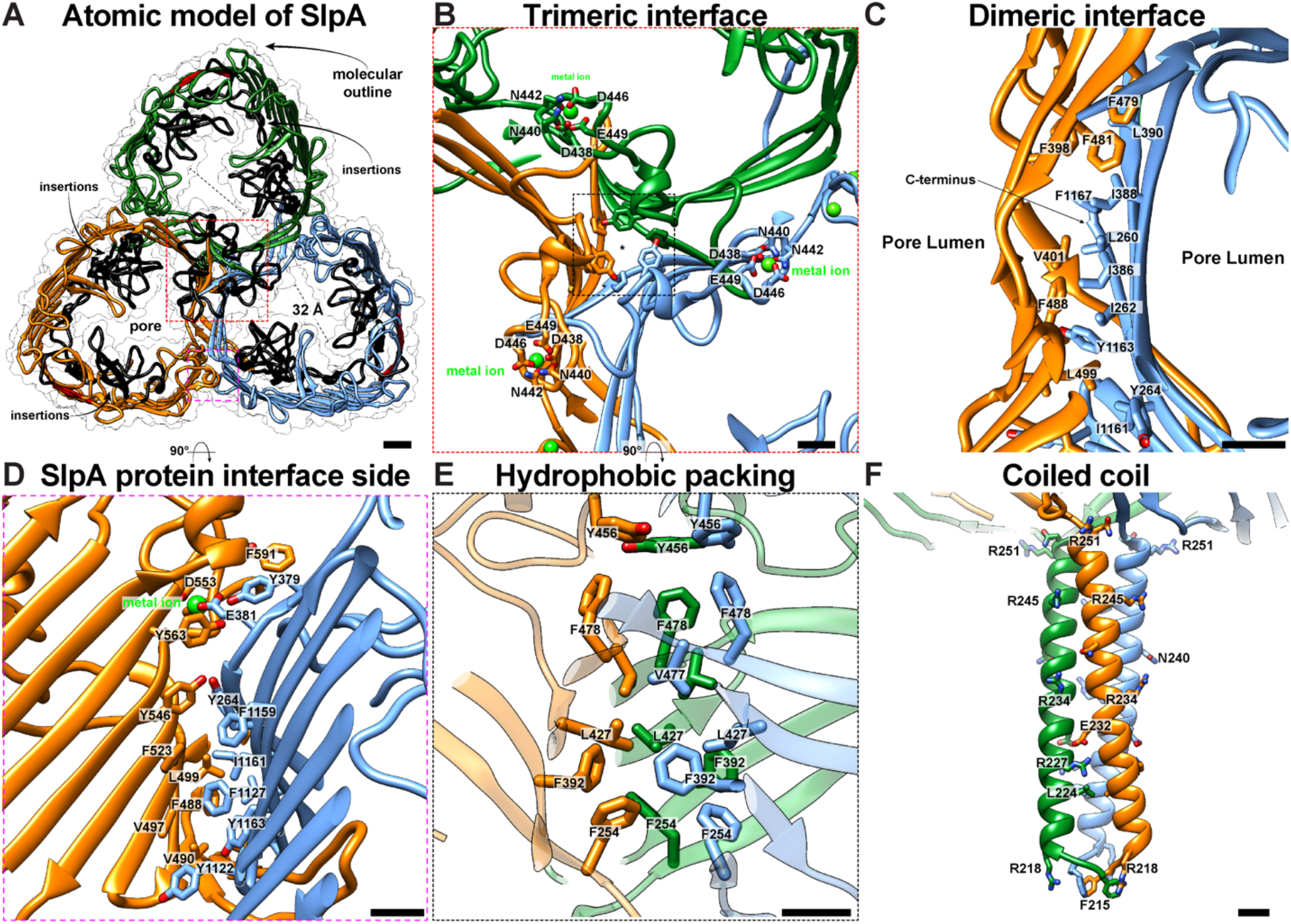
Atomic structure of trimeric outer membrane porin. (A) The refined atomic model of the trimeric SlpA protein shown as a ribbon diagram. The pore is blocked by several insertions (black ribbons). (B) Close-up view of the central trimeric interface is shown where a typical insertion including metal binding sites is found. (C-D) The dimeric SlpA:SlpA interface is lined by hydrophobic residues and stabilized by a metal binding site (see Figure S4). (E) The central trimeric interface is stabilized by hydrophobic packing of aromatic residues as shown in an orthogonal, magnified view of panel B. (G) Close-up view of the end of the coiled-coil segment is shown. Scale bars: A) 10 Å; B-F) 5 Å.

The protein:protein interface of the SlpA β-barrel consists mainly of stacked β-sheets from apposing barrels (Figures 2C-D and S4). These sheets contain a hydrophobic patch made up of residues L260, I262, Y264, I386, I388, L390, and F392 stacking onto V401, F479, F481, F488 and L499 from the next subunit (Figure 2C-D). At the trimeric interface (C3 axis), another set of hydrophobic residues F254, F392, L427, V477, and F478 stabilize the complex (Figure 2E).

There is clear density in the map for residues 215-253 that make up the coiled-coil segment connected with the OMBB (Figures 2F and S1). The coiled coil consists of a highly conserved, prominent salt bridge between E232 and R227 (Figures 2F and S4). Residue R245 points away from the axis of the coiled coil and is bound to a poorly resolved density for a previously uncharacterized protein rich in β-strands (Figure S5). We were able to ascertain the identity of this protein using the map density combined with structural modelling (Uniprot DR_0644); however, due to the weak density, atomic model refinement of this newly identified protein was not performed. Homologs of this accessory protein are found in many other Deinococcales, but are absent in Thermales. The residues of the SlpA coiled coil prior to residue 215 are less well resolved in our cryo-EM map (Figure S1), but diffuse density for the N-terminal part of the protein extends well beyond the well-resolved part of the coiled coil (Figures 1D and S5A). This extended arrangement of the coiled coil supports SlpA’s role in bridging the OM, where the β-barrel (residues 254-1167) is situated, and the PG, which is expected to bind to the N-terminal SLH domain (predicted residues 29-92, Figure S2).

### Structural modelling of the periplasmic part of the *D. radiodurans* SlpA

Next, we used the power of the recently developed, state-of-the-art structure prediction method AlphaFold-Multimer (33), which has been shown to yield fairly accurate atomic models of homo- and hetero-meric complexes (34), to model the periplasmic part of *D. radiodurans* SlpA. This part of the sequence (residues 20-252), comprising the SLH domain and a section of the coiled coil, was poorly resolved in our map. The model yielded by AlphaFold-Multimer had high per-residue confidence (pLDDT) and low Predicted Aligned Error (PAE) values, both of which are indicators for high accuracy of the model (Figure S6). In fact, the part of *D. radiodurans* homotrimeric coiled-coil segment resolved in our cryo-EM map (residues 215-254) and the corresponding part in the AlphaFold-Multimer model showed remarkable similarity, superimposing with a root mean square deviation (RMSD) of ∼0.39 Å over all Cα atoms. The complete model of the N-terminal part of *D. radiodurans* SlpA shows that the length of the homotrimeric coiled-coil segment is approximately 28 nm, which is in good agreement with our measurements from cryo-EM (Figure S5, ∼29 nm). The coiled-coil segment exhibits two β-layers (Figure S2), which are triangular supersecondary structural elements formed in trimeric coiled coils to compensate for local strains resulting from the insertion of two or six amino acids into the canonical heptad repeats(35).

Next, we analyzed the periplasmic segments of several other SlpA proteins at an organizational level (Figure 3). The length of the coiled-coil segment of SlpA is comparably long in other Deinococcales and is even longer in Thermales (Figures 3 and S7), supporting a role for SlpA as an OM-PG connector or spacer. Moving towards the IM, the coiled-coil segment is connected to the SLH domain via a short, disordered linker in *D. radiodurans* (Figure S2). The SLH domain of SlpA, like other previously characterized SLH domains, is also predicted to form a trimer. The SLH domain is highly conserved among Deinococcus-Thermus SlpA proteins (Figure S8), with an average pairwise sequence identity of greater than 60% and conserved sequence motifs (W, residue 14; GVILG, residues 54-57; and TRYE, residues 70-73 in *D. radiodurans* SlpA) characterized to be important for interactions with PG-linked SCWPs in other SLH domains (25, 36, 37). This agrees with our cryo-EM data, suggesting that the *D. radiodurans* SLH domain connects the OMBB to the PG at the N-terminal end of the coiled coil.

**Figure 3.**
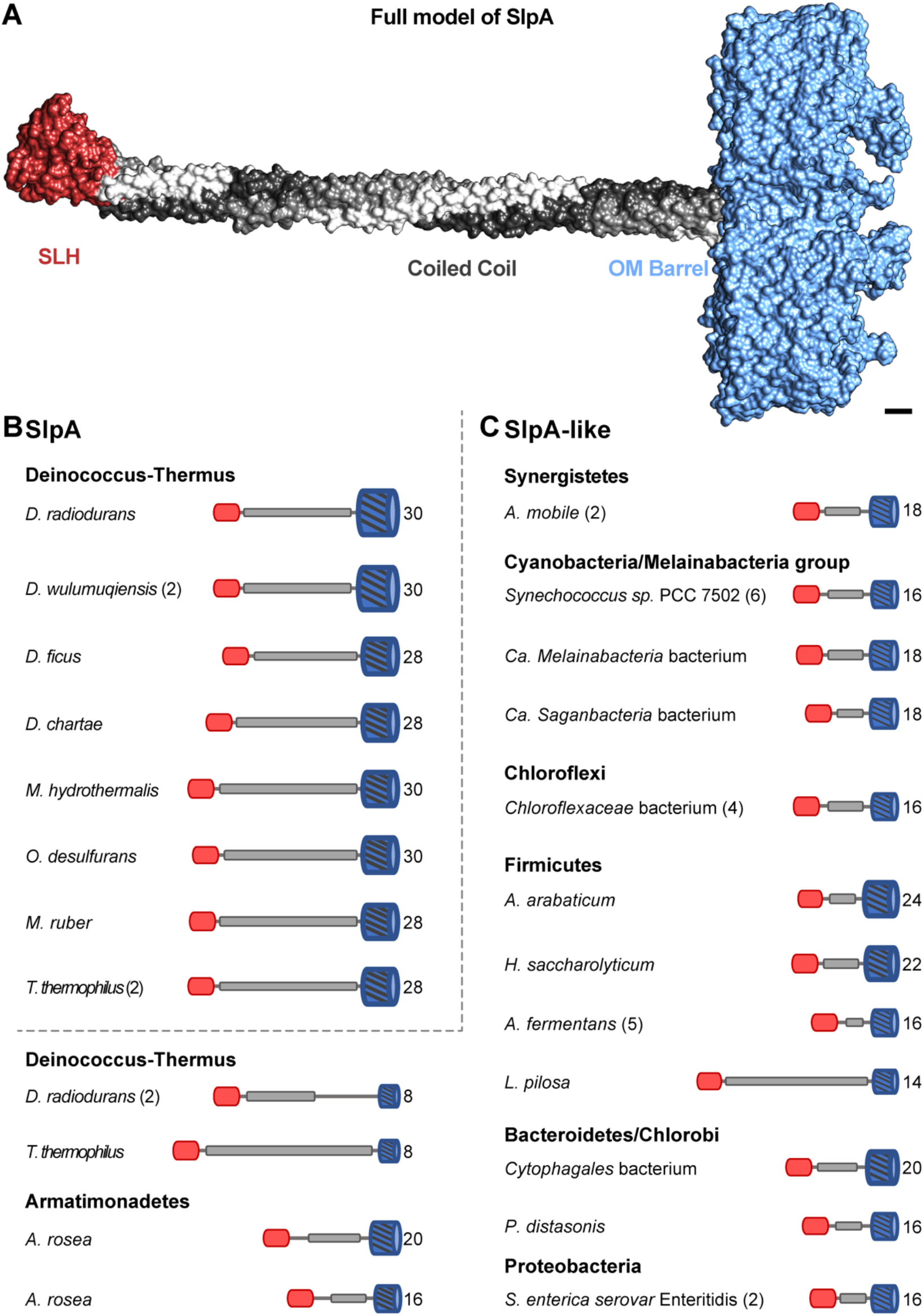
Structural modelling of SlpA-like proteins reveals common organizational principles of OM-PG connectors. (A) Combined cryo-EM and AlphaFold model of SlpA from *D. radiodurans.* Scale bar: 10 Å. (B) The domain organization of representative SlpA and SlpA-like proteins from bacteria of several Gram-negative phyla are shown. While they exhibit a shared tripartite organization, comprising an N-terminal SLH domain, a central coiled-coiled segment, and a C-terminal OMBB, the length of the coiled-coil segment and the number of β-strands (indicated on the right end of the cartoons) in the OMBB is quite varied. Some organisms, such as *D. wulumuqiensis* and *T. thermophilus*, contain two or more paralogs (indicated within rounded brackets). Accession details for the shown proteins are provided in Table S2.

### Several OMBB proteins in deep-branching Gram-negative bacteria contain coiled coils connected to SLH domains

To investigate the presence of other SlpA-like proteins in *D. radiodurans,* we searched for all OMBB-containing proteins in its genome using the predictive power of HHpred (38) and AlphaFold (34). In addition to SlpA, we found 19 further OMBB-containing proteins, with three predicted to form large, 38-stranded β-barrels (Table S3). Curiously, similarly to SlpA, two of these 19 OMBB proteins also contain a central coiled-coil segment and an N-terminal SLH domain possessing residues important for binding PG-linked SCWPs (Figure S8). However, unlike SlpA, they comprise an 8-stranded β-barrel that is reminiscent of the β-barrel of the outer membrane protein A (OmpA) (39), which is an OM-PG tether found in some Gram-negative bacteria such as *Escherichia coli*. Homologs of these two SlpA-like *D. radiodurans* proteins are widespread in the Deinoccocus-Thermus phylum, suggesting that they, like SlpA, might also be involved in connecting the OM to the inner cell envelope. The presence of several OMBBs as well as the recently described PilQ secretin complex that traverses both membranes (40), suggests that even though SlpA is a highly abundant molecule in the *D. radiodurans* cell envelope, it cannot fully tile the OM and is probably not an integral part of the S-layer. We, however, cannot rule out that it may play a sub-stoichiometric, minor role in anchoring the HPI protein.

We next investigated if SlpA-like proteins are also present in other deep-branching Gram-negative bacterial lineages, because their presence could represent an ancestral mechanism for tethering the OM to the inner cell envelope. To this end, we searched and detected a widespread occurrence of SlpA-like proteins in the deep-branching phyla Synergistetes, Cyanobacteria, Candidatus Melainabacteria, Armatimonadetes, and Bacteroidetes (Figure 3). Furthermore, we also found a widespread occurrence of SlpA-like proteins in the Gram-negative lineages Halanaerobiales, Negativicutes, and Limnochordia of the largely Gram-positive phylum Firmicutes as well as a sparse occurrence in Chloroflexi, Chlorobi, and Proteobacteria. These SlpA-like proteins are frequently annotated as iron uptake porin, carbohydrate porin, S-layer protein, S-layer homology domain-containing protein, or hypothetical protein in protein sequence databases, but we could not find any experimentally characterized representatives. Although SlpA-like proteins exhibit a tripartite domain organization as SlpA of *D. radiodurans* and possess sequence motifs important for interactions with PG-linked SCWPs in their SLH domain, they contain OMBBs of varying sizes and much shorter coiled-coil segments (Figure 3), which is expected, given that the periplasmic space in *D. radiodurans* is substantially thicker compared to most other Gram-negative bacteria. We predict that these SlpA-like proteins probably also form homotrimeric complexes that link the OM to the inner envelope.

### Model of the *D. radiodurans* cell envelope

To relate our atomic structural and bioinformatic data with the native cell envelope, we next collected electron cryotomograms of whole *D. radiodurans* cells and envelopes of partly lysed cells (Figures S9). In line with previous reports (9, 11), we observed a cell envelope with two membranes, a large periplasmic space of 121 ± 4 nm and a thick PG layer. As expected from our structural results, we observed a fuzzy density corresponding to the start of the wide PG layer at a distance of 30 ± 3 nm from the OM, in agreement with the length of periplasmic coiled-coil segment of SlpA observed in our atomic model (28 nm) and class averages (29 nm, Figure S9). Outside the OM, we observed that the S-layer did not uniformly coat the entire surface of cells, but was rather found as large patches on the cell surface (Figure S9), 18 ± 1 nm away from the OM.

Taken together, we report an updated model of the *D. radiodurans* cell surface (Figure 4). We suggest that while SlpA does not tile the OM, due to the presence of several other OMBBs in the *D. radiodurans* genome. The observed patches of S-layer could be held in a sub-stoichiometric manner by OMBBs of SlpA, which are present in abundance in the OM, where they are connected through coiled-coil segments to the PG layers via SLH domains.

**Figure 4.**
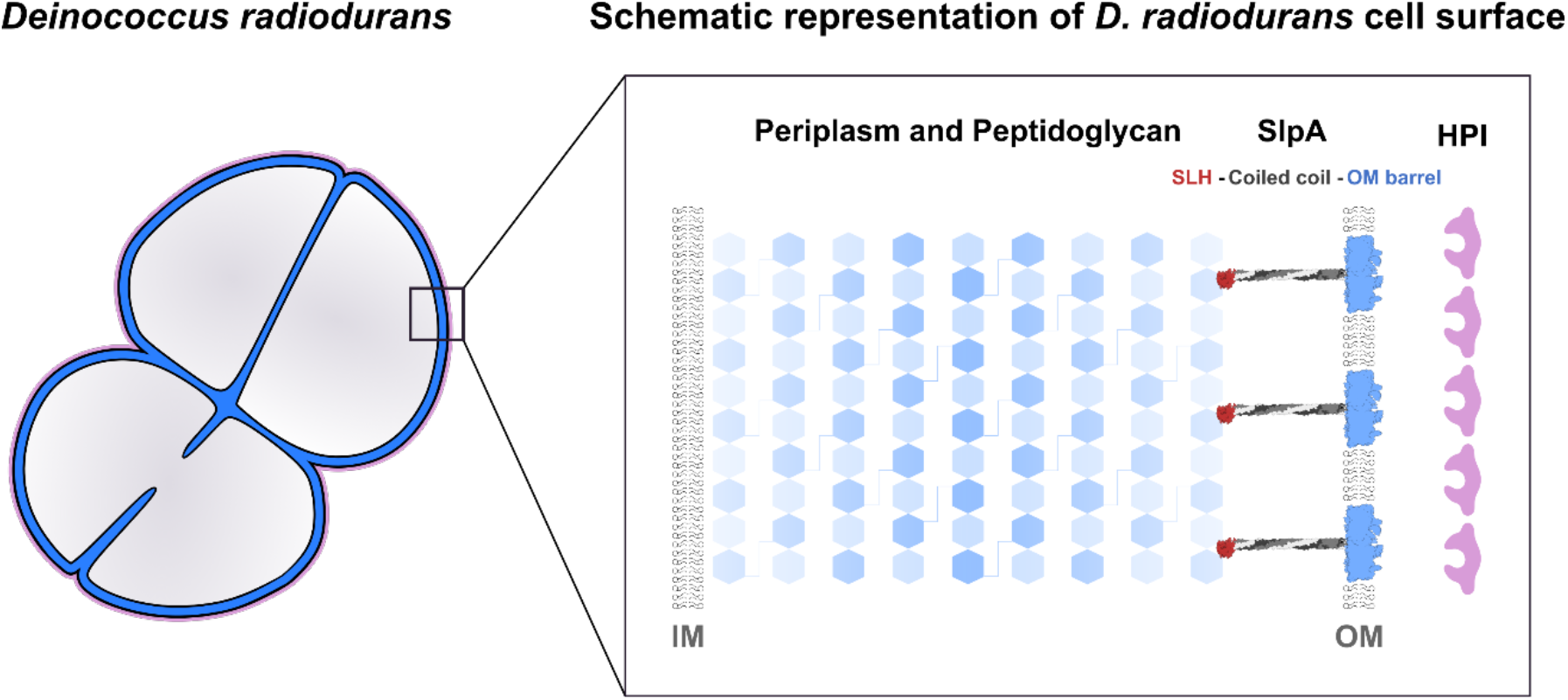
Model of the *D. radiodurans* cell envelope. Schematic model of the *D. radiodurans* cell envelope shows how SlpA connects the OM to the PG layer via long coiled coils and an N-terminal SLH domain, placing data from previous studies into context, and providing a structural framework for understanding the cell envelope of deep-branching Gram-negative bacteria.

## Discussion

In this study, we present structural data to resolve a longstanding conundrum about the role of the OM protein SlpA in organizing the cell surface of *D. radiodurans*. While initial studies suggested that the S-layer of *D. radiodurans* is built by the HPI protein, more recent studies have proposed that it is formed of multiple proteins, including HPI and SlpA. Our results indicate that SlpA cannot fully tile the OM and that it is not a fundamental component of the S-layer, but that it connects the OM to the PG layer by forming extended homotrimers. SlpA might play a minor, sub-stoichiometric role in anchoring the HPI protein, although this role of SlpA has not been demonstrated. SlpA exhibits a tripartite organization, comprising an OMBB trimer embedded in the OM, a long coiled-coil stalk, and an SLH domain trimer, typically found in Gram-positive SLPs (24). Combining our atomic structures and bioinformatic results with tomography of native cell envelopes, we report an updated model for the complex, multi-layered cell envelope of *D. radiodurans* (Figure 4), which will serve as a structural framework for understanding the cell surface of similar deep-branching bacteria with atypical envelopes.

Furthermore, we show that SlpA-like proteins, frequently containing OMBBs of varying sizes and coil-coiled segments of varying lengths, are widespread in the Deinococcus-Thermus phylum as well as in several phyla of deep-branching Gram-negative bacteria, suggesting that they represent an ancestral mechanism for stabilizing the cell envelope by connecting the OM to the PG layer. In some Proteobacteria, such as *E. coli*, *Coxiella burnetii*, *Pseudomonas aeruginosa*, and *Legionella pneumophila*, highly abundant OM proteins that form covalent or non-covalent connections between the OM and the PG layer have been shown to be important for the stabilization and the spacing of the OM with respect to the IM. Such proteins include Braun’s lipoprotein (Lpp) (41, 42), PG-associated lipoprotein (Pal) (43), and the OMBB proteins OmpA (39, 43), OprF (44, 45), and BbpA (46). However, OM-PG connectors remain poorly characterized in most other phyla of Gram-negative bacteria. The physiological role of SlpA-like proteins we detected in this study remain unknown currently, but we speculate that they may be involved in maintaining the integrity of the OM in several phyla of Gram-negative bacteria.

We also expect that the multi-domain architecture of SlpA is crucial for its role as an organizational spacer in the cell envelope. To illustrate this, in the same manner as *D. radiodurans*, the deep-branching hyperthermophilic bacterium *Thermotoga maritima* also exhibits an unusual cell envelope that is thought to be stabilized by two equally abundant OM proteins, Ompα and Ompβ, which resemble SlpA (47, 48). While Ompα has been characterized to be a rod-shaped spacer in electron micrographs, Ompβ forms triangular, porin-like assemblies in the OM. Like SlpA, Ompα contains an N-terminal SLH domain and a long coiled-coil segment, which has been predicted to be 45 nm in length; however, instead of an OMBB, the C-terminal end of Ompα contains a transmembrane helix that anchors it to the OM. The identity of Ompβ has not been established experimentally yet, but an OMBB, encoded by a gene that occurs adjacent to the gene encoding Ompα in an operon in *T. maritima* and some closely related organisms, is likely to be Ompβ (49), and this OMBB is predicted by AlphaFold to contain 22 β-strands. We speculate that, like SlpA, Ompα and Ompβ associate to form a homotrimeric complex, a scenario that would be consistent with both the OMBB and the coiled-coil part of SlpA-like proteins being important for acting as a spacer, critical for organising the cell envelope. Moreover, while *T. maritima* contains two further paralogs of Ompα, which have also been implicated to play a role in the organization of its cell envelope (50), orthologs of Ompα and Ompβ are widespread in the phylum Thermotogae (49).

Questions of whether monoderm or diderm bacteria came first, and how and when the transition between them occurred are major open questions in evolutionary biology (51–53). The widespread occurrence of the SLH domain in monoderm and diderm bacterial proteins, including SLPs and SlpA-like proteins, suggests that the SLH domain was established as a PG-binding domain very early in the evolution of bacteria. Furthermore, given the widespread occurrence of SlpA-like proteins in diderm bacteria and the role of SlpA in organizing the OM of *D. radiodurans*, it appears plausible that an ancestral SlpA-like protein was already present and functioned as an OM-PG connector in the common ancestor of diderm bacteria. It is therefore tempting to speculate that irrespective of whether monoderm or diderm bacteria were first, SLH domain-containing proteins may have been involved in allowing the loss or gain of the OM during the transition between monoderm and diderm bacteria, and it would be fascinating to explore this possibility moving forward.

## Materials and Methods

### SlpA protein purification

*Deinococcus radiodurans* cells from ATCC (ATCC BAA816) were grown in modified tryptone-glucose-yeast extract (TGY) medium supplemented with 5 µM MnCl_2_(54). For protein purification of wild-type SlpA, 4 L of modified TGY medium were inoculated 1:25 with a late-log phase pre-culture, and cells were grown overnight with shaking at 30 °C. Cells were harvested by centrifugation (5,000 relative centrifugal force (rcf), 4 °C, 30 minutes), and the cell pellet was resuspended in 50 mL lysis buffer (100 mM Tris/HCl pH 8.0, 150 mM NaCl, 5 mM MgCl_2_, 50 µg/mL DNAseI, 1 U/mL benzonase (SigmaAldrich), 1x cOmplete protease inhibitor (Roche)) per 1 L cell pellet. Cells were lysed by passing the suspension five times through a homogenizer at 22,500 pounds per square inch (psi), and unlysed cells were removed by centrifugation (2,000 rcf, 4 °C, 15 minutes). Remaining cell debris was isolated by centrifugation (48,000 rcf, 4 °C, 30 minutes). To degrade PG, the pellet was resuspended in 40 mL lysozyme buffer (100 mM Tris/HCl pH 8.0, 500 µg/mL lysozyme, 1 x cOmplete Inhibitor) and incubated on a rotary wheel for 16 hours at 4 °C. The remaining insoluble fraction was pellet by centrifugation (48,000 rcf, 4 °C, 30 minutes), washed three times with 37.5 mL wash buffer (100 mM Tris/HCl pH 8.0, 150 mM NaCl) and separated centrifugation after each step (48,000 rcf, 4 °C, 30 minutes). Membrane proteins in the final washed pellet were resuspended in 40 mL buffer (20 mM Tris/HCl pH 8.0) and extracted with detergent by adding drop-wise a 10% (w/v) stock solution of n-dodecyl β-D-maltoside (DDM, Anatrace) to a final concentration of 1.3% (w/v). The protein suspension was next incubated on a rotary wheel for 3 hours at 4 °C and non-solubilized material was removed by centrifugation (30,000 rcf, 4 °C, 30 minutes). The protein solution was then loaded onto an equilibrated 5 mL HiTrap-Q columns (GE Healthcare) using an ÄKTA pure 25 system (GE Healthcare) and unbound protein was washed away with 50 mL binding buffer (20 mM Tris/HCl pH 8.0, 0.05% (w/v) DDM). Bound protein was eluted with an increasing gradient of 75 mL elution buffer (20 mM Tris/HCl pH 8.0, 0.05% (w/v) DDM, 1 M NaCl). Fractions containing SlpA were pooled, concentrated using a 30 kDa molecular weight cut-off (MWCO) Ultra Centrifugal tube (Amicon) and loaded to a Superose 6-Increase 10/300 GL column (GE Healthcare) equilibrated with 20 mM HEPES/NaOH pH 7.5, 150 mM NaCl, 0.02 % (w/v) DDM. Protein was eluted in the same buffer, and fractions containing SlpA were collected, concentrated (Amicon 30 kDa MWCO) to 200 µL, and then dialyzed against 100 mL SEC buffer (20 mM HEPES/NaOH pH 7.5, 150 mM NaCl, 0.02 % (w/v) DDM) for 2 hours with a 10 kDA MWCO cutoff. For cryo-EM grid preparation the final protein solution was then concentrated to 4.45 mg/mL and immediately used for cryo-EM grid preparation. Purified SlpA was kept at 4 °C, reloaded onto a Superose 6 Increase 10/300 GL column (GE Healthcare) and analyzed by SDS-PAGE, which showed minimal degradation upon prolonged storage. Chromatograms and SDS-PAGE gel images were visualized with MATLAB (MathWorks) and Fiji(55), respectively.

### Cryo-EM sample preparation

For cryo-EM grid preparation, 2.5 µL of 4.45 mg/mL SlpA sample or sonicated (5 s, 15 mA amplitude) late-log cultures *D. radiodurans* culture were applied to a freshly glow discharged Quantifoil R2/2 Cu/Rh 200 mesh grid, adsorbed for 10 s, blotted for 5 s and plunge-frozen into liquid ethane in a Vitrobot Mark IV (ThermoFisher), while the blotting chamber was maintained at 100% humidity at 10 °C. For cryo-ET, 10 nm protein-A gold (CMC Utrecht) was additionally added to the samples immediately prior to grid preparation.

### Cryo-EM data collection and single particle analysis

Data collection: Single-particle cryo-EM data were collected on a Titan Krios G3 microscope (ThermoFisher) operating at 300 kV fitted with a Quantum energy filter (slit width 20 eV) and a K3 direct electron detector (Gatan) with a sampling pixel size of 0.546 Å running in counting super-resolution mode. For the SlpA specimen, a total of 2,294 movies were collected in two sessions with a dose rate of 2.98 e^-^/pixel/s on the camera level. The sample was subjected to 4.8 s of exposure, during which a total dose of 47.909 e^-^/Å^2^ respectively was applied, and 40 frames were recorded per movie (see Table S1).

Image processing: Movies were clustered into optics groups based on the XML meta-data of the data-collection software EPU (ThermoFisher) using a k-means algorithm implemented in EPU_group_AFIS (https://github.com/DustinMorado/EPU_group_AFIS). Imported movies were motion-corrected, dose weighted, and Fourier cropped (2x) with MotionCor2 (56) implemented in RELION3.1 (57). Contrast transfer functions (CTFs) of the resulting motion-corrected micrographs were estimated using CTFFIND4 (58). Initially, micrographs were denoised using TOPAZ (59) using the UNET neural network and 2893 particles were manually picked. Particle coordinates were used to train TOPAZ picker (60) in 5x downsampled micrographs with the neural network architecture ResNet8 and picked particles were extracted in 4x downsampled 128 x 128 boxes and classified using reference-free 2D classification inside RELION3.1. An initial subset of 76,119 particles were then used to re-train TOPAZ, followed by another round of particle extraction and reference-free 2D classification. Particles belonging to class averages with high-resolution features were combined, and duplicate particles within 100 Å where removed, and an initial model was generated with 4x downsampled particles in 128 x 128 boxes using the SGD-algorithm within RELION3.1. The initial reference was aligned to the C3 symmetry axis and the merged particle subset was re-extracted in 512 x 512 boxes and subjected to a focused 3D auto refinement on the central porin and the first heptad of the coiled coil using the scaled and lowpass filtered output from the symmetry aligned starting model. Per-particle defocus, anisotropy magnification and higher-order aberrations (57) were refined inside RELION-3.1, followed by signal subtraction of the detergent micelle and another round of focused 3D auto refinement. The reconstruction was further improved by Bayesian particle polishing (61), and a focused 3D-classification without refinement of the poses. The final map was obtained from 122,412 particles and post-processed using a soft mask focused on the central trimer including the first heptad of the coil coiled yielding a global resolution of 3.25 Å according to the gold standard Fourier shell correlation criterion of 0.143 (62). Cryo-EM single-particle data statistics are summarized in Table S1.

### Cryo-ET data collection, tomogram segmentation and subtomogram averaging

Data collection: For tomographic data collection, the SerialEM software (63) was used as described previously (64). Tomographic data collection of cellular specimens was performed on the same Titan Krios microscope as above using the Quantum energy filter (slit width 20 eV) and the K3 direct electron detector running in counting mode. Tilt series (with a defocus range of −8 to −11 µm were collected between ±60° in a dose symmetric scheme (65) with a 2° tilt increment. A total dose of 121 e^-^/Å^2^ with a dose-rate of 10.523 e^-^/px/s was applied over the entire series, and image data were sampled at a pixel size of 3.468 Å.

Image processing: Tilt series alignment using gold fiducials and tomogram generation was performed in IMOD (66). Tensor voting based membrane detection was performed with TomosegmemTV (67), refined and visualized in Chimera (68) and ChimeraX (69). Distances between inner membrane, peptidoglycan layer, outer membrane and S-layer were determined at multiple positions along the cell surface throughout the tomogram. Subtomogram averaging analysis of the *D. radiodurans* cell surface was performed using previously described methods (70, 71), also previously applied to Gram-negative bacterial cell surfaces (72).

### Model building and refinement

The carbon backbone of the SlpA protein was manually traced through a single subunit of the cryo-EM density using Coot (73). The atomic model was subjected to several rounds of refinement using REFMAC5 (74) inside the CCP-EM software suite (75) and PHENIX (76), followed by manually rebuilding in Coot (73) and interactive refinement using ISOLDE (77) inside UCSF ChimeraX (69). Model validation was performed in PHENIX and CCP-EM, and data visualization was performed in UCSF Chimera (68) and ChimeraX.

### Bioinformatic analysis

A structural model of the periplasmic, homotrimeric segment (residues 20-252) of *D. radiodurans* SlpA was built using an installation of AlphaFold-Multimer (33) at the Max Planck Computing and Data Facility in Garching. The prediction was carried out in default settings, and the model (ranked_0.pdb) with the highest confidence was picked for further use (Figure S6). Homologs of *D. radiodurans* SlpA in the Deinococcus-Thermus phylum were detected using the NCBI BLAST Web server in default settings (78). To detect SlpA-like proteins in Gram-negative bacteria, we used a three-step approach. First, we searched the non-redundant protein sequence database at NCBI for homologs of the SLH domain of *D. radiodurans* SlpA; the search was restricted to Gram-negative phyla of bacteria. Next, we inspected the obtained sequences for the presence of a central coiled-coil segment using PCOILS (79) and a C-terminal OMMB using HHpred (80) searches against the ECOD (81) profile Hidden Markov Model (HMM) database. Finally, the three-dimensional structures of some representative SlpA-like proteins (Table S2) were predicted using AlphaFold (34). To detect OMBB proteins in the proteome of *D. radiodurans*, we searched its profile HMM database with HHpred in the MPI Bioinformatics Toolkit(38). The searches were seeded with sequences of OMBBs from the ECOD X group ‘Outer membrane meander beta-barrels’. Next, to analyze the domain composition and the number of β-strands in the barrel, we built structural models of the obtained matches using AlphaFold (Table S2). Multiple sequence alignments of the SLH (Figure S8), coiled-coil (Figure S7), and OMMB (Figure S3) domains were calculated using PROMALS3D (82), and were subsequently curated manually based on AlphaFold models.

## Acknowledgments

T.A.M.B. is a recipient of a Sir Henry Dale Fellowship, jointly funded by the Wellcome Trust and the Royal Society (202231/Z/16/Z). T.A.M.B. would like to thank the Vallee Research Foundation, the European Molecular Biology Organization, the Leverhulme Trust and the Lister Institute for Preventative Medicine for support. The authors would like to thank Jan Löwe and Danguole Kureisaite-Ciziene for providing the *D. radiodurans* strain. V.A. would like to thank Andrei Lupas for continued support. This work was partly supported by institutional funds of the Max Planck Society.

## Supplementary Information for

**Other supplementary materials for this manuscript include the following:**

Movies S1

### Supplementary Figures

**Figure S1.**
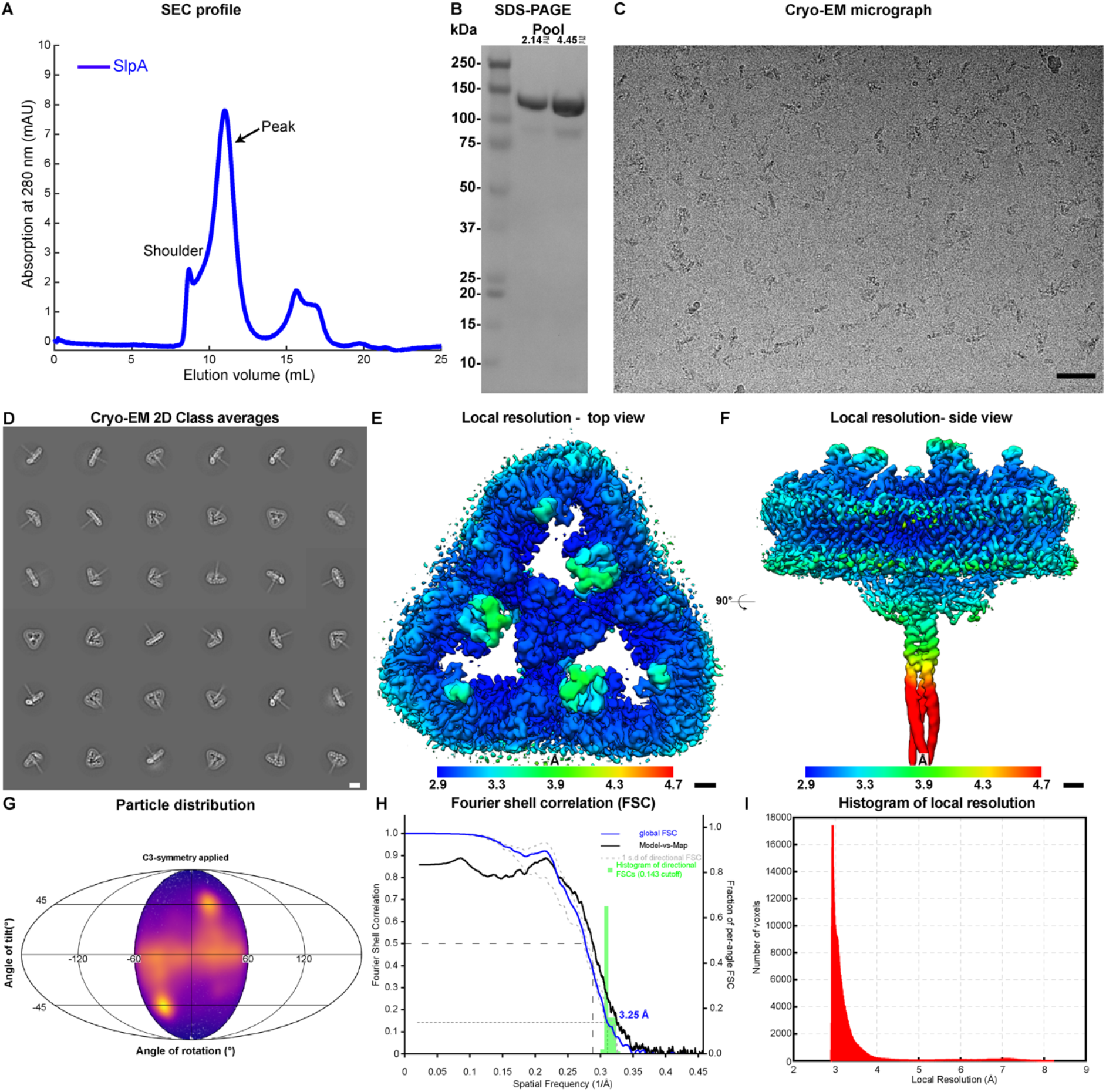
Cryo-EM structure of the SlpA porin of *D. radiodurans*. (A) Analytical size-exclusion chromatography (SEC) profile and (B) SDS-PAGE analysis of the final purified SlpA protein from the native source. (C) Cryo-EM image of purified SlpA protein flash frozen in liquid ethane (density black). (D) 2D Class averages of SlpA particles picked using TOPAZ (1) and classified inside RELION-3.1 (2) while ignoring CTF-correction until the first peak (density white). (E-F) Local resolution estimated in RELION, plotted into the map, shown in two orthogonal orientations. (G) Angular distribution of the particles in the data set. (H) 3D Fourier shell correlation between two random halves of the data (FSC) (3) and Model vs Map FSC. Histogram binning size is set to 5. (I) Histogram of local resolutions in voxels of the cryo-EM map. Scale bars: C) 500 Å; D) 100 Å; E-F) 10 Å.

**Figure S2.**
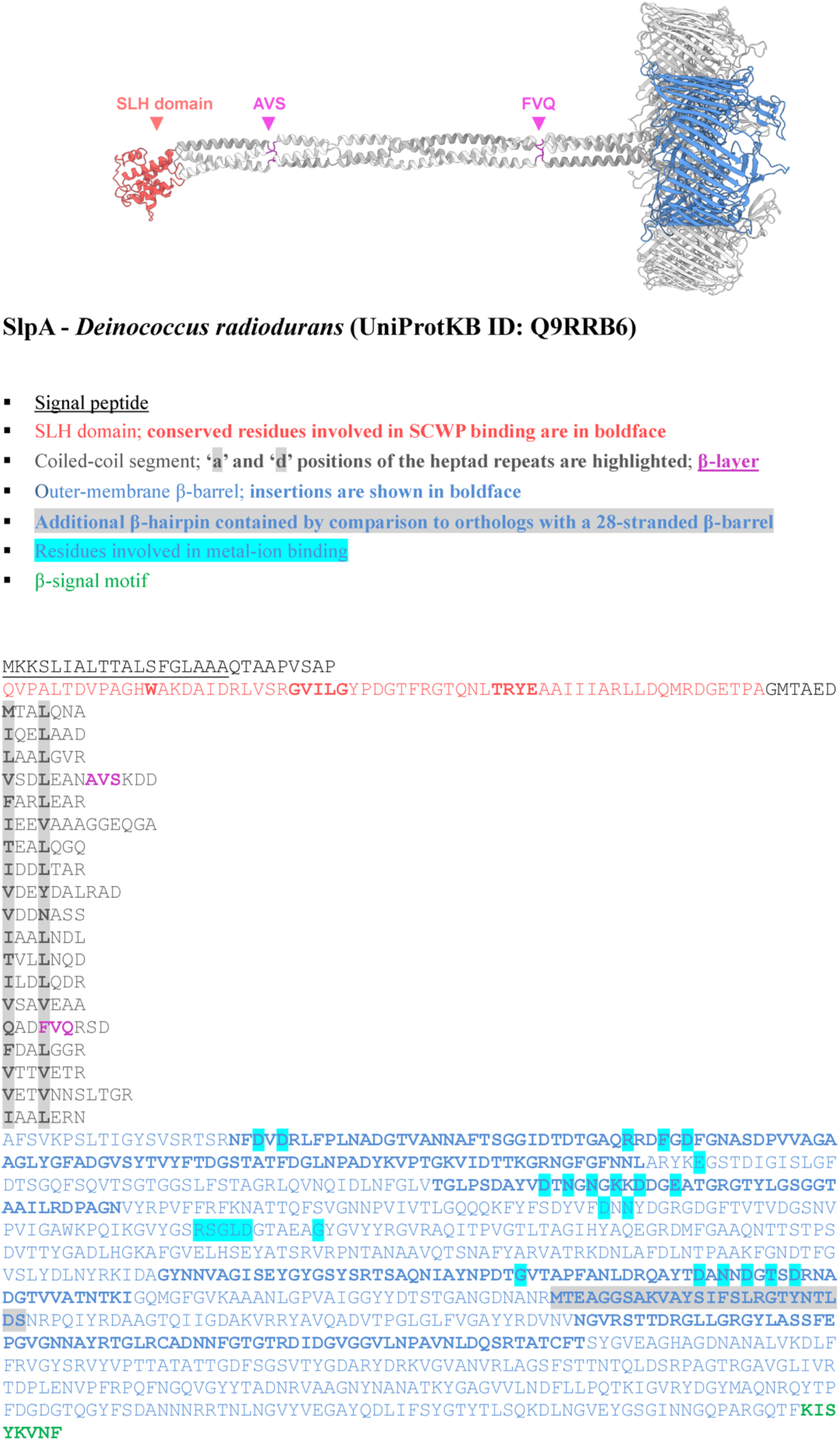
Sequence annotation of the *D. radiodurans* SlpA protein. A full structural model of *D. radiodurans* SlpA obtained by combining the cryo-EM and AlphaFold models is shown in cartoon representation; the two β-layers are colored in magenta (upper). The boundaries of the SLH domain, the coiled-coil segment, and the OMBB as well all other sequence features, as detailed in the rectangular box, are marked in the sequence of SlpA (lower).

**Figure S3.**
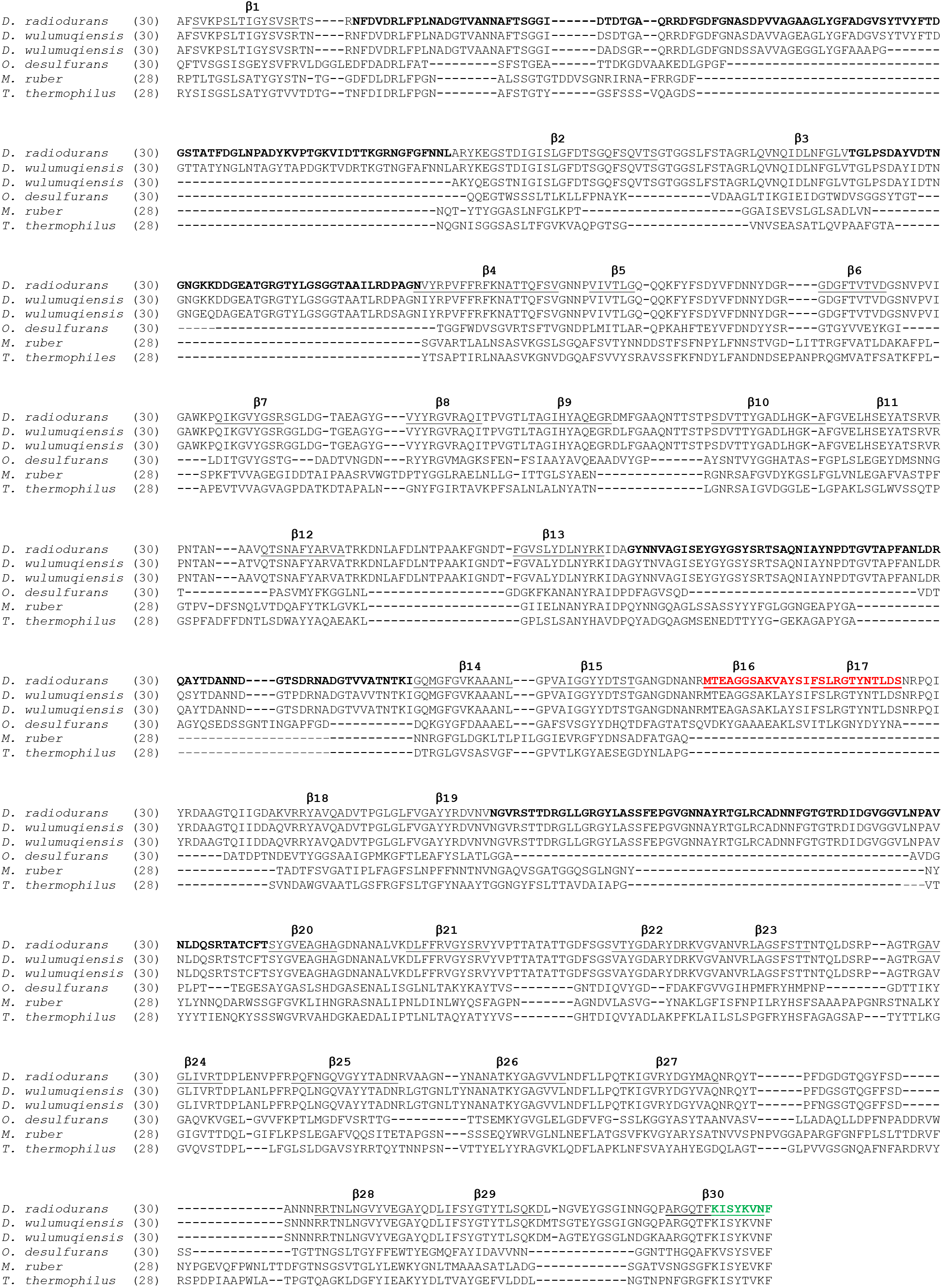
Multiple sequence alignment of the OMBB region of representative SlpA proteins. Residues part of β-strands are underlined in the sequence of *D. radiodurans*. Large insertions in *D. radiodurans* OMBB are shown in boldface, the extra β-hairpin is colored red, and the β-signal motif (BAM insertion signal) is colored green. The number of β-strands contained in the individual OMBBs are indicated with rounded brackets. Accession details for the shown sequences are provided in Table S2.

**Figure S4.**
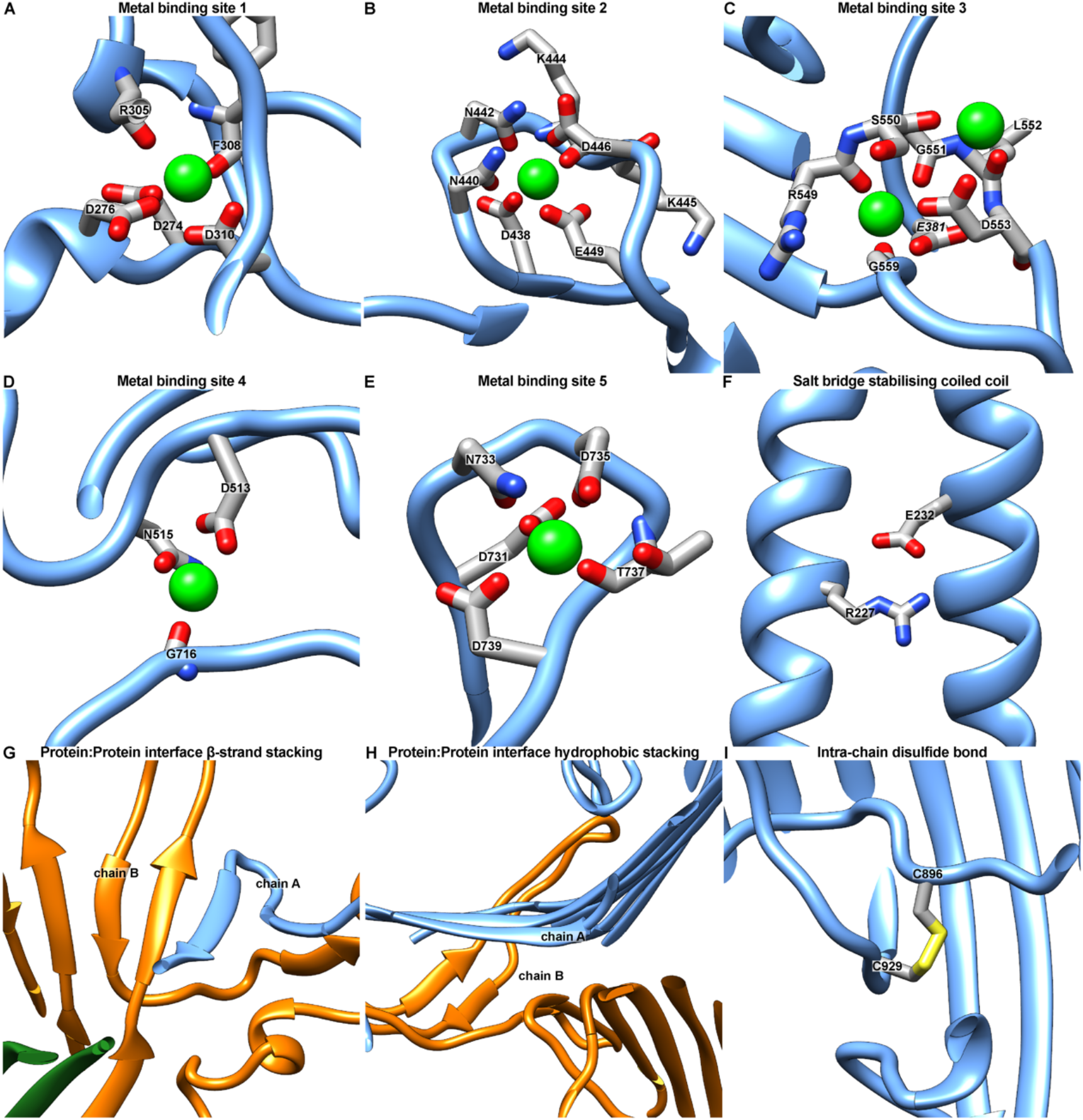
Close-up views of the SlpA OMBB model. (A-E) Close up views of putative metal ion (green) binding sites in SlpA, depicted as ribbon and stick diagram. In all metal binding sites, putative metal ions are coordinated by carboxyl group of aspartate and glutamate residues as well as the carbonyl oxygen of asparagine side chains or the main chain peptide bond. (F) A prominent salt bridge between the E232 and R227 of a second SlpA subunit stabilizes the coiled-coil region. (G) Close to the central symmetry axis the trimeric conformation is stabilized by hydrogen bonding via β-strand stacking. (H) The β-sheets of two opposing outer membrane barrels are stabilized through hydrophobic stacking (see also Figure 2C,D). (I) The primary sequence of SlpA contains two cysteine (896, 929) residues which form an intra-chain disulfide bond within the plug of the OMBB.

**Figure S5.**
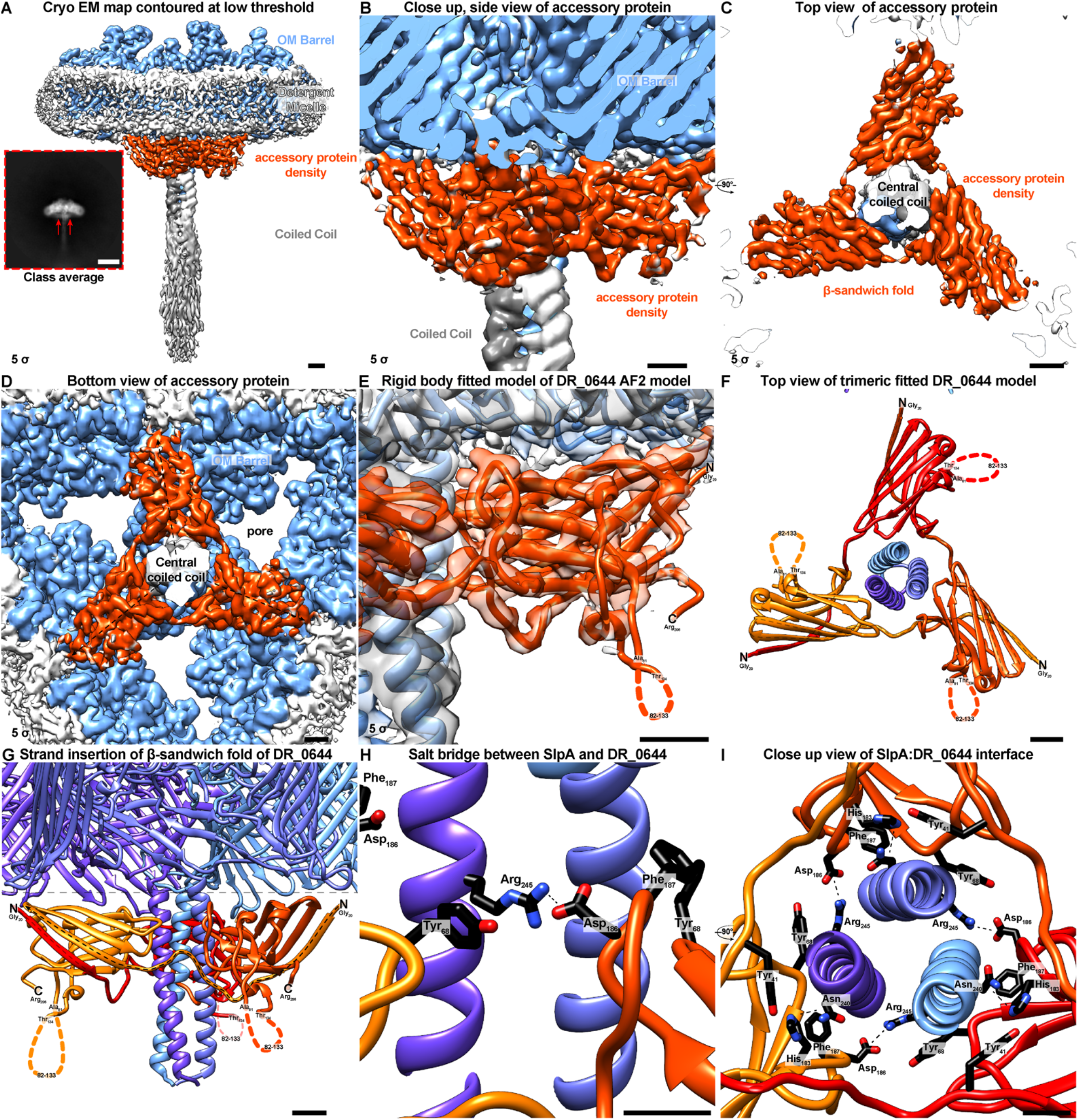
Additional protein density bound to SlpA. (A) Density map of SlpA OMBB trimer at a contour level of 5 σ shown in light blue embedded in a detergent micelle (white). An additional protein density (orange) is partially resolved, present in a subclass of particles seen in 2D class averages (Inset, Scale bar Inset: 100 Å). (B-D) Close up view of the additional density seen from the side B), top C) and bottom D). The accessory protein adopts a trimeric β-sandwich fold which is aligned with the OMBB without occluding the pore region. (E) Rigid body fit of the Alphafold2 (4) (AF2) model of the uncharacterized protein DR_0644, which has been previously identified to be associated with SlpA (5), into the extra density not explained by SlpA. The model of the protein fits exceptionally well from the mature N-terminus (N, Gly_20_) to the C-terminus (C, Arg_206_), with the exception of an unstructured loop (82-133) which is in an agreement with its low confidence of the predicted local-distance difference test (pLDDT) as measured by AF2. (F-G) The β-sandwich fold of the trimeric DR_0644 (red, orange red, and orange ribbons) is completed by strand insertion of the first β-strand of the next clockwise oriented subunit, as seen from the top F) and side G). (H-I) The interface of the SlpA:DR_0644 is stabilized by a prominent salt bridge between SlpA R245 and DR_0644 D186 and further protein:protein interactions such as hydrogen bonding between SlpA N240 and DR_0644 H183. Scale bars: A-G) 10 Å; H-I) 5 Å.

**Figure S6.**
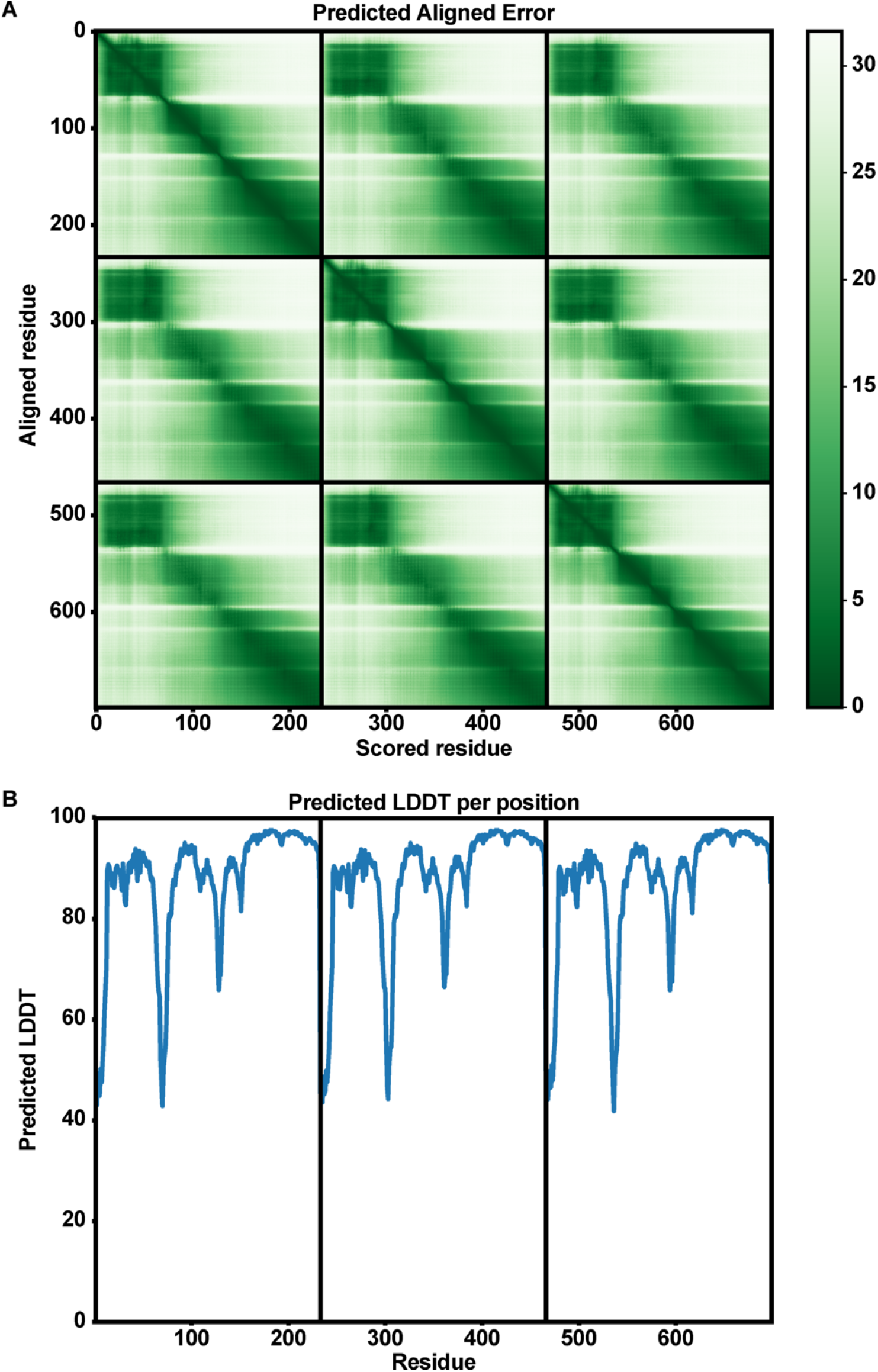
AlphaFold modelling of the SlpA periplasmic segment. The homotrimeric periplasmic part of SlpA was modelled using AlphaFold-Multimer. The Predicted Aligned Error (PAE, upper) and per-residue confidence (pLDDT, lower) plots for the model with the best confidence is shown. Low PAE and high pLDDT values are indicators for high accuracy of the model yielded by AlphaFold. The subplots in each panel correspond to one SlpA monomer.

**Figure S7.**
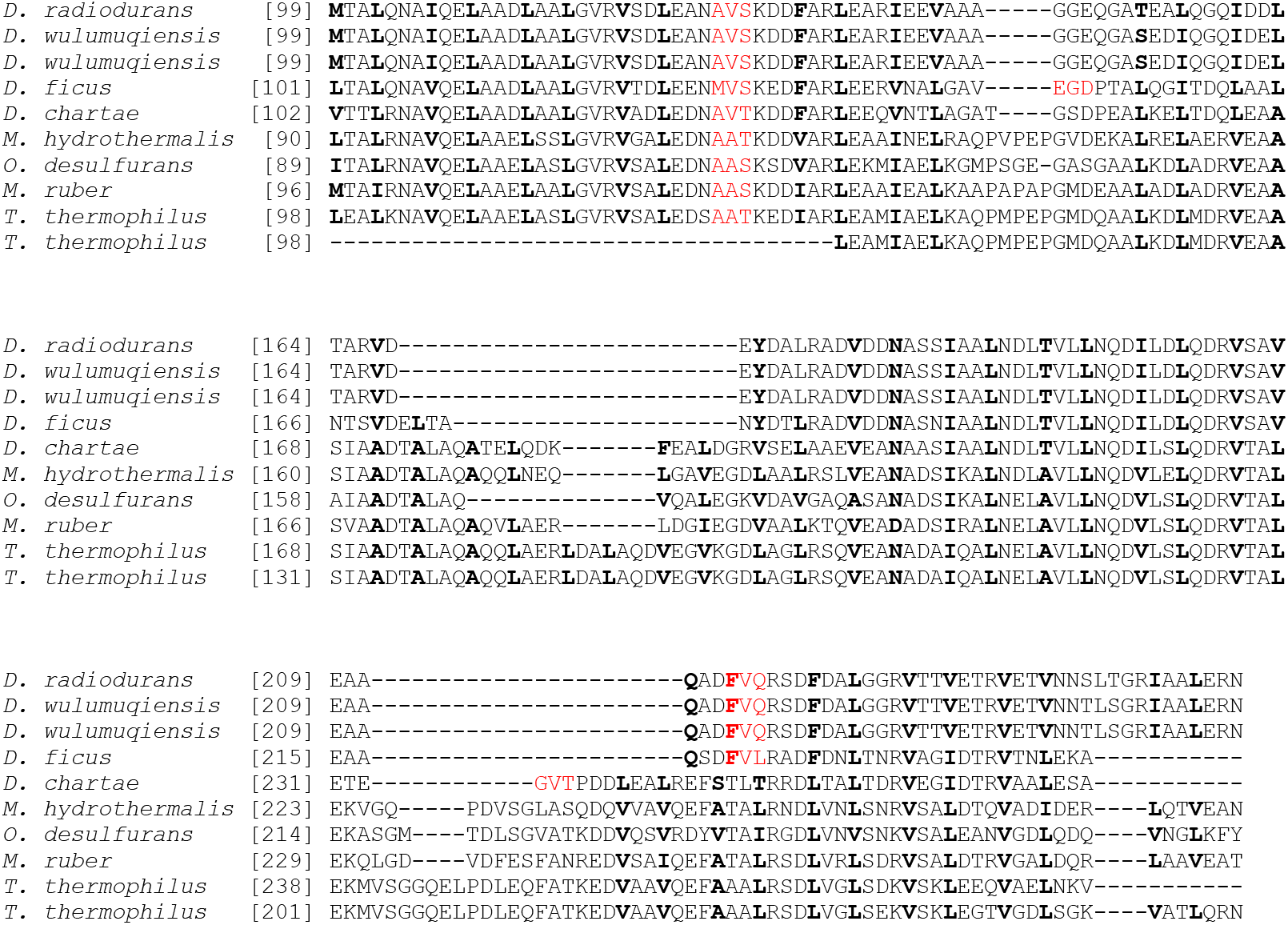
Multiple sequence alignment of the coiled-coil region of representative SlpA proteins. The core-forming hydrophobic positions (‘a’ and ‘d’ positions of the heptad repeats in canonical coiled coils) are shown in boldface in each sequence. Residues involved in forming β-layers are coloured red. Accession details for the shown sequences are provided in **Table S2**.

**Figure S8.**
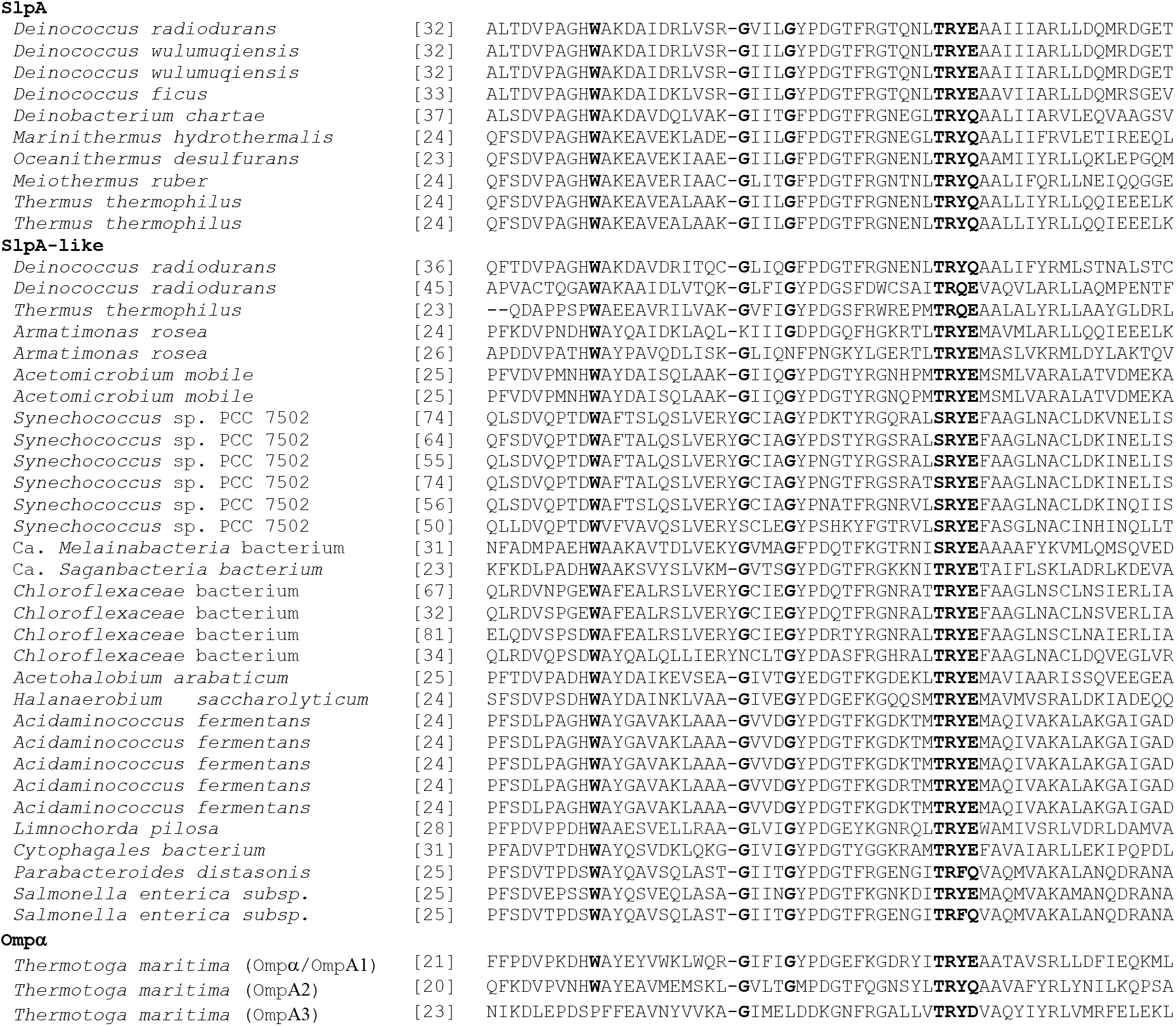
Sequence alignment of SLH domains from SlpA-like proteins. Residues important for binding PG-linked SCWPs are shown in boldface, and the start position of the SLH domain is indicated in brackets. Accession details for the shown sequences are provided in Table S2.

**Figure S9.**
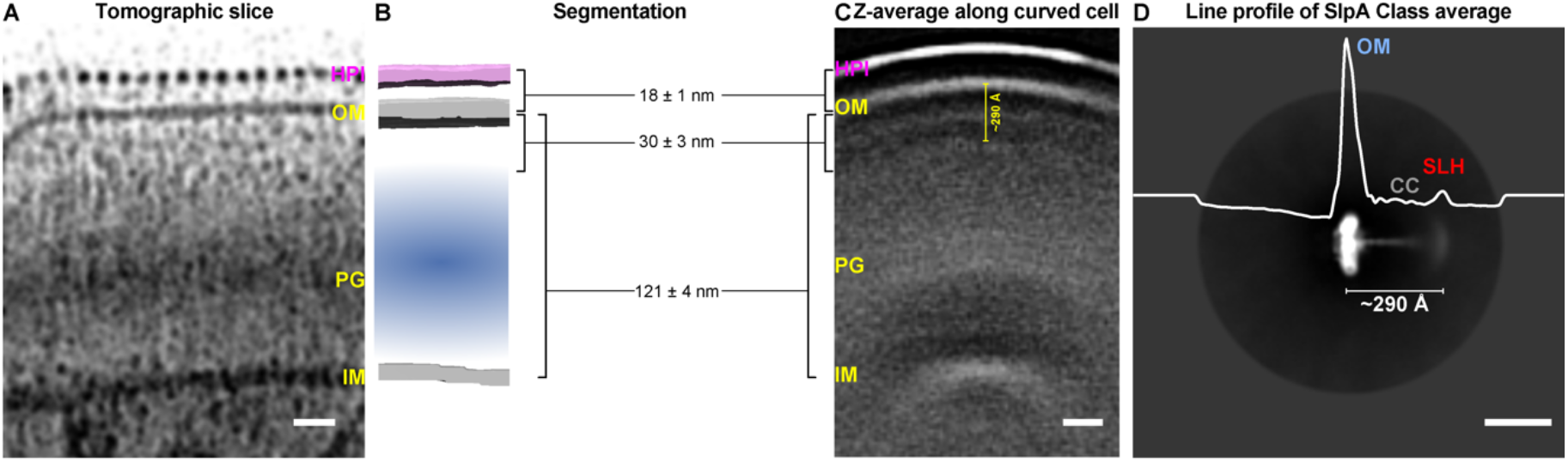
Cryo-ET and cryo-EM of *D. radiodurans* cell envelopes. (A) Tomographic slice through the outer envelope of a partially lysed *D. radiodurans* cell. The cell envelope can be divided into four prominent regions, as listed from the extracellular space to the cytosol: A repetitive S-layer containing the HPI protein, the OM, a thick layer of PG, and the IM, which is in agreement with previous studies (6). (B) Segmentation of the tomographic volume. (C) Z-average of subtomograms extracted along the curved surface of *D. radiodurans* highlights the most prominent cell surface features of the complex envelope of *D. radiodurans*. (D) Line profile of 2D cryo-EM class average of SlpA with the same distance of the OMBB to the S-layer homology (SLH) domain (marked). Scale bar: A,C,D) 200 Å.

### Supplementary Tables

**Table S1:** Cryo-EM data collection, refinement and validation statistics.

|  |  |
| --- | --- |
|  | #Drad_SlpA |
|  | (EMDB-xxxx) |
|  | (PDB xxxx) |
| **Data collection and processing** |  |
| Microscope | Titan Krios G3 |
| Magnification | 81,000 |
| Voltage (kV) | 300 |
| Total Electron exposure (e-/Å<sup>2</sup>) | 47.909 |
| Detector | K3 (Gatan) |
| Slit width (eV) | 20 |
| Defocus range (μm) | -1 to -2.5 |
| Acquisition Mode | Super-resolution |
| Pixel size (Å) | 0.546 |
| AFIS<sup>&</sup> Mode | Yes |
| Micrographs collected | 2,294 |
| Micrographs used | 2,294 |
| **Data processing** |  |
| Software reconstruction | RELION3.1 (2) |
| Software picking | TOPAZ (1) |
| Initial particle images (no.) | 223,878 |
| Final particle images (no.) | 122,412 |
| Rescaled Box-size Class2D (px) | 128 x 128 |
| Rescaled Box-size Class3D (px) | 512 x 512 x 512 |
| Final Box-size (px) | 512 x 512 x 512 |
| Pixel size final reconstruction (Å) | 1.092 |
| Symmetry imposed | C3 |
| Map resolution (Å) | 3.25 |
| FSC threshold | 0.143 |
| Map resolution range (Å) | 2.9 – 6.7 |
| Map sharpening *B* factor (Å<sup>2</sup>) | -46.8126 |
| 3DFSC sphericity<sup>#</sup> | 0.985 |
| **Model Refinement** |  |
| Initial model used (PDB code) | None |
| Software | PHENIX (7) |
| Model resolution (Å) | 3.3 |
| FSC threshold | 0.5 |
| Model composition |  |
| Non-hydrogen atoms | 21,504 |
| Protein residues | 2,859 |
| Ions | 18 |
| *B* factors (Å<sup>2</sup>) |  |
| Protein | 51.67 |
| Ligand | 58.42 |
| R.m.s. deviations |  |
| Bond lengths (Å) | 0.002 |
| Bond angles (°) | 0.472 |
| Validation |  |
| MolProbity score | 1.29 |
| Clashscore | 3.61 |
| Poor rotamers (%) | 0.00 |
| Cβ outliers (%) | 0.00 |
| CABLAM outliers (%) | 2.14 |
| Ramachandran plot |  |
| Favored (%) | 97.27 |
| Allowed (%) | 2.73 |
| Disallowed (%) | 0.00 |
| Rama-Z (Z-score, RSMD) |  |
| whole (N= 2853) | -1.01 (0.16) |
| helix (N= 183) | -0.65 (0.42) |
| sheet (N= 1236) | -0.39 (0.15) |
| loop (N= 1434) | -0.83 (0.16) |

& AFIS: Aberration Free Imaging Shift Mode.

**Table S2:** Accession information for SlpA and SlpA-like proteins; several bacteria contain multiple homologs.

| Organism | NCBI/UniProt accession | Barrel size |
| --- | --- | --- |
| **SlpA** |  |  |
| *Deinococcus radiodurans* | WP_010889202 | 30-stranded |
| *Deinococcus wulumuqiensis* | WP_017871655 | 30-stranded |
| *Deinococcus wulumuqiensis* | WP_114672325 | 30-stranded |
| *Deinococcus ficus* | WP_191241882 | 28-stranded |
| *Deinobacterium chartae* | WP_183988817 | 28-stranded |
| *Marinithermus hydrothermalis* | WP_013702952 | 30-stranded |
| *Oceanithermus desulfurans* | WP_147148761 | 30-stranded |
| *Meiothermus ruber* | WP_013015232 | 28-stranded |
| *Thermus thermophilus* | WP_011228992 | 28-stranded |
| *Thermus thermophilus* | P35830 | 28-stranded |
| **SlpA-like** |  |  |
| *Deinococcus radiodurans* | WP_010887767 | 8-stranded |
| *Deinococcus radiodurans* | WP_027480303 | 8-stranded |
| *Thermus thermophilus* | WP_011228967 | 8-stranded |
| *Armatimonas rosea* | WP_184200374 | 20-stranded |
| *Armatimonas rosea* | WP_184200710 | 16-stranded |
| *Acetomicrobium mobile* | WP_014807739 | 18-stranded |
| *Acetomicrobium mobile* | WP_014806458 | 18-stranded |
| *Synechococcus* sp. PCC 7502 | WP_015167070 | 16-stranded |
| *Synechococcus* sp. PCC 7502 | WP_015168280 | 16-stranded |
| *Synechococcus* sp. PCC 7502 | WP_015168863 | 16-stranded |
| *Synechococcus* sp. PCC 7502 | WP_015168481 | 16-stranded |
| *Synechococcus* sp. PCC 7502 | WP_041429169 | 16-stranded |
| *Synechococcus* sp. PCC 7502 | AFY74914 | 16-stranded |
| *Ca. Melainabacteria bacterium* | PKL78650 | 18-stranded |
| *Ca. Saganbacteria bacterium* | MBI5699473 | 18-stranded |
| *Chloroflexaceae* bacterium | NJL83094 | 16-stranded |
| *Chloroflexaceae* bacterium | NJL82168 | 16-stranded |
| *Chloroflexaceae* bacterium | NJL82038 | 16-stranded |
| *Chloroflexaceae* bacterium | NJL82082 | 16-stranded |
| *Acetohalobium arabaticum* | WP_013279101 | 24-stranded |
| *Halanaerobium saccharolyticum* | WP_133516444 | 22-stranded |
| *Acidaminococcus fermentans* | WP_106787916 | 16-stranded |
| *Acidaminococcus fermentans* | WP_012939007 | 16-stranded |
| *Acidaminococcus fermentans* | WP_012939010 | 16-stranded |
| *Acidaminococcus fermentans* | WP_012939008 | 16-stranded |
| *Acidaminococcus fermentans* | WP_012939009 | 16-stranded |
| *Limnochorda pilosa* | WP_068139859 | 14-stranded |
| *Cytophagales* bacterium | MBC8103537 | 20-stranded |
| *Parabacteroides distasonis* | WP_172733356 | 16-stranded |
| *Salmonella enterica* subsp. | EBY4570518 | 16-stranded |
| *Salmonella enterica* subsp. | EBY4570338 | 16-stranded |
| **Ompα and Ompβ** |  |  |
| *Thermotoga maritima* (Ompα/OmpA1) | Q01969 | - |
| *Thermotoga maritima* (OmpA2) | Q9X252 | - |
| *Thermotoga maritima* (OmpA3) | Q9WZH0 | - |
| *Thermotoga maritima* (OMPβ) | WP_004081490 | 22-stranded |

**Table S3.** OMBB proteins in *D. radiodurans*; proteins for which we are unsure about the number of β-strands are indicated by (?).

| NCBI accession | Barrel size | Remarks |
| --- | --- | --- |
| Q9RRB6 | 30-stranded | SlpA |
| AAF11946 | 38-stranded | Contains N-terminal immunoglobulin-like (invasin D3) domains |
| UID71982 | 38-stranded | Contains N-terminal immunoglobulin-like (invasin D3) domains |
| UID69687 | 38-stranded | Contains N-terminal immunoglobulin-like (invasin D3) domains |
| AAF11400 | 8-stranded | No additional domains |
| ANC71147 | 8-stranded | Contains a C-terminal domain with a cystatitin-like (NFT2-like) fold |
| ANC71478 | 18-stranded | No additional domains |
| AAF10759 | 8-stranded | No additional domains |
| AAF10697 | 8-stranded | SlpA-type |
| AAF10549 | 8-stranded | No additional domains |
| UID69670 | 8-stranded | No additional domains |
| AAF10255 | 8-stranded | No additional domains |
| AAF10153 | 38-stranded (?) | LPS-assembly protein LptD |
| AAF09960 | 16-stranded | Outer membrane assembly factor BamA |
| ANC72597 | 8-stranded | SlpA-type |
| QIP33551 | 24-stranded | PapC usher |
| ANC70518 | 12-stranded (?) | No additional domains |
| AAF09880 | 12-stranded (?) | No additional domains |
| AAF11062 | 8-stranded | No additional domains |
| AAF10975 | 8-stranded | No additional domains |

### Supplementary Movie Legend

**Movie S1. Cryo-EM reconstruction of SlpA from *D. radiodurans***

The 3.25 Å cryo-EM reconstruction is shown with the atomic model (ribbon diagram) built into the density (isosurface).

